# Prediction of the sex-determination gene in tunas (*Thunnus* fishes)

**DOI:** 10.1101/2021.06.15.448492

**Authors:** Yoji Nakamura, Kentaro Higuchi, Kazunori Kumon, Motoshige Yasuike, Toshinori Takashi, Koichiro Gen, Atushi Fujiwara

**Affiliations:** Bioinformatics and Biosciences Division, Fisheries Stock Assessment Center, Fisheries Resources Institute, Japan Fisheries Research and Education Agency, 2-12-4 Fuku-ura, Kanazawa, Yokohama, Kanagawa 236-8648, Japan; Tuna Aquaculture Division, Aquaculture Research Department, Fisheries Technology Institute, Japan Fisheries Research and Education Agency, 1551-8 Taira-machi, Nagasaki, 851-2213, Japan; Amami Field Station, Tuna Aquaculture Division, Aquaculture Research Department, Fisheries Technology Institute, Japan Fisheries Research and Education Agency, 955-5 Hyousakiyamahara, Setouchi, Kagoshima 894-2414, Japan; Aquatic Breeding Division, Aquaculture Research Department, Fisheries Technology Institute, Japan Fisheries Research and Education Agency, 422-1 Nakatsuhamaura, Minami-ise, Mie, 516-0193, Japan

**Keywords:** bluefin tuna, sex determination, gene duplication, estrogen sulfotransferase

## Abstract

Fish species have a variety of sex determination systems. Tunas (genus *Thunnus*) have an XY genetic sex-determination system. However, the Y chromosome or responsible locus has not yet been identified in males. In a previous study, a female genome of Pacific bluefin tuna (*T. orientalis*) was sequenced, and candidates for sex-associated DNA polymorphisms were identified by a genome-wide association study using resequencing data. In the present study, we sequenced a male genome of Pacific bluefin tuna by long-read and linked-read sequencing technologies, and explored male-specific loci through a comparison with the female genome. As a result, we found a unique region carrying the male-specific haplotype, where a homolog of estrogen sulfotransferase gene was predicted to be encoded. The genome-wide mapping of previously resequenced data indicated that, among the functionally annotated genes, only this gene, named *sult1st6y*, was paternally inherited in the males of Pacific bluefin tuna. We reviewed the RNA-seq data of southern bluefin tuna (*T. maccoyii*) in the public database and found that *sult1st6y* of southern bluefin tuna was expressed in all male testes, but absent or suppressed in the female ovary. Since estrogen sulfotransferase is responsible for the inactivation of estrogens, it is reasonable to assume that the expression of *sult1st6y* in gonad cells may inhibit female development, thereby inducing the individuals to become males. Thus, our results raise a promising hypothesis that *sult1st6y* is the sex-determination gene in *Thunnus* fishes, or at least functions at a crucial point in the sex-differentiation cascade.

## Introduction

Fish species have a substantial variety of sex-determination systems compared to other animals, ranging from environmentally to genetically modulated ones. In the environmental sex determination system, individual sex depends mainly on nongenetic factors such as temperature, growth rate, and density, while in the genetic system, specific genes dominantly control sex differentiation. Regarding sexuality, it is estimated that more than 98% of fish are gonochoristic (Pandian, 2011). Therefore, in most fish, some responsible loci should exist biasedly between males and females or differentially affect gonadal phenotypes, even if the sex is primarily determined by environmental factors. In heterogametic (XY) males or heterogametic (ZW) females, sex chromosomes may be cytologically evident markers as carrying the sex determination loci, but they are often homomorphic in fishes (Devlin and Nagahama, 2002), undistinguishable from autosomes by direct microscopic observation. In other words, fish sex chromosomes are not highly differentiated, implying that turnovers may have often occurred over the course of evolution (Pennell et al., 2018). Recently, sex determination genes have been reported in several fish species, including *DMY/Dmrt1* in medaka (Matsuda et al., 2002; Nanda et al., 2002) and Chinese tongue sole (Cui et al., 2017), *Amhr2* in fugu (Kamiya et al., 2012), *sdY* in rainbow trout (Yano et al., 2012), *Hsd17b1* in yellowtail (Koyama et al., 2019), and *amh* in Patagonian pejerrey (Hattori et al., 2012) and northern pike (Pan et al., 2019). These studies suggest that a master gene for sex determination in fish can be easily replaced by another gene, which may be followed by the above-mentioned turnover of the sex chromosome. In particular, the cases in fugu and yellowtail are prominent examples, where only single nucleotide substitutions are critical for sex determination, suggesting that the replacements of master gene occurred only recently in these lineages. Thus, the sex determination genes of fish are often different for each lineage and are not always identified by deductive approaches from model organisms (that is, case studies are required to identify the genes).

Regarding tunas, the fish of the genus *Thunnus* South 1845, it is estimated that Pacific bluefin tuna (*Thunnus orientalis*, PBT) has an XY genetic sex determination system (Agawa et al., 2015). However, the chromosome or locus responsible has not yet been identified. One reason for this is that no distinguishable sex chromosomes were observed in the cytological analysis. It has been reported that little tuna, *Euthynnus affinis*, may have a ZW sex determination system (Yazawa et al., 2019). These observations suggest that the sex determination gene of *Thunnus* species has only recently emerged, and therefore, that the sex chromosomes are not highly divergent. Another reason is the technical difficulty in handling: since tunas are large and difficult fish to culture, *in vivo* experiments for identifying sex-associated loci are costly and time-consuming. On the other hand, because tunas have a high market value worldwide, tuna target fishing and aquaculture have been conducted in many countries. In aquaculture, the control of the sex ratio in tanks or ocean nets is very important for breeding, as keeping more females than males leads to an increased production of fertilized eggs. Since tunas do not show any clear sexual dimorphism, the direct observation of the gonad has been the main technique for sex identification. However, this is a destructive inspection and is not applicable for selectively collecting females or removing males in tanks. Therefore, molecular markers associated with the sex of tuna have been of interest as an alternative for rapid sex identification. In particular, DNA markers have been considered promising, where a small portion of tissue sections, such as the fin or skin, are sufficient for PCR assays. A DNA marker associated with PBT sex was reported in 2015 (Agawa et al., 2015). The marker, Md6, was developed from the DNA samples of cultured PBT individuals and showed an association with sex in the aquaculture population. Regarding the genomic approach, the PBT male genome data were open (Nakamura et al., 2013), but the scaffold sequences were too fragmented for a genome-wide screening of sex-associated loci. Recently, transcriptomics of southern bluefin tuna (*T. maccoyii*) have been conducted, and the differentially expressed genes between male and female gonads (testis and ovary) were examined (Bar et al., 2016). However, it should be noted that a differentially expressed gene between the testis and ovary is not always the sex determination gene, but may be a downstream gene regulated by the master gene.

In 2019, Suda et al. sequenced the genome of female PBT, compared the assembled scaffold with that of the male genome, and explored sex-associated regions from the scaffold sequences by resequencing 31 PBT individuals (15 males and 16 females) (Suda et al., 2019). They performed a genome-wide association (GWA) study focusing on sex-associated single nucleotide variants (SNVs) between males and females, and identified several regions carrying male-specific alleles. Finally, they developed three PCR primer pairs for sex identification, namely scaf64_3724604_F/R, scaf64_3726411_F/R, and scaf64_3724591_F/R, based on male-specific heterozygous SNVs and short indels detected in the 6.5-kb region of female scaffold 64 (F64). The PCR primers were designed so that the amplification patterns were different between PBT males and females: scaf64_3724604_F/R and scaf64_3726411_F/R were amplifiable in males (the sizes were 113 bp and 143 bp, respectively) but not in females, and scaf64_3724591_F/R was amplified in both males and females, but the sizes were different (142 bp and 149 bp in males and only 149 bp in females). PCR amplification using 115 individuals (56 males and 59 females) yielded 100% accuracy of sex identification, implying that the PBT sex-determination gene may exist around the 6.5-kb region in F64. They predicted some genes in F64 and discussed the possible involvement in the sex determination of PBT. However, promising genes, such as those related to the cascade, were not identified. The potential problem in the previous study was that they found male-specific heterozygous polymorphisms in multiple scaffolds. Since most sex determinants are often attributed to a single locus, they suspected that multiple candidates occurred due to assembly errors.

In this study, we improved the male genome and examined the sex-associated regions using previously published data. We have found a sex-associated gene that is paternally inherited and is involved in estrogen inactivation. The current results provide the most promising candidate for the sex determination gene of tuna.

## Materials and Methods

### Genome sequencing and assembly

A three-year-old male F2 individual of Pacific bluefin tuna, which was cultured at the Seikai National Fisheries Research Institute (currently Fisheries Technology Institute), was collected for genome sequencing. Frozen heart tissue was prepared from the specimen, and DNA was extracted using a DNeasy Blood & Tissue Kit (Qiagen, Venlo, Netherlands). Then, an SMRT-bell template library was prepared following the manufacturer’s protocol and sequenced using the Pacific Biosciences (PacBio) Sequel platform (P6C4 chemistry) (Pacific Biosciences, Inc. CA). From the same tissue sample, the DNA was extracted also using the standard phenol-chloroform protocol. A Chromium 10X library was prepared using Chromium Genome Library Kit v2, Genome Gel Bead Kit, Genome Chip Kit v2, and i7 Multiplex Kit (10X Genomics, CA) following the manufacturer’s protocol, and the linked reads were sequenced using the Illumina NovaSeq 6000 platform (Illumina, Inc. CA). The PacBio reads were first assembled by HGAP4 (SMRT Link v5.1.0.26412) (Chin et al., 2013). The contigs obtained were corrected using NovaSeq linked-reads by Pilon v1.23 (Walker et al., 2014) and Tigmint v1.1.2 (Jackman et al., 2018), and finally scaffolded by ARKS v1.0.2 (Coombe et al., 2018) in combination with LINKS v1.8.5 (Warren et al., 2015). The scaffold sequences obtained were submitted to the DNA Data Bank of Japan under the accession nos.

BOUD01000001-BOUD01000948. To examine the phased genomic regions, the linked reads were assembled using Supernova2 (v2.1.1) (Weisenfeld et al., 2017). The former assembly (HGAP4 + ARKS) was assessed by Benchmarking Universal Single-Copy Orthologs (BUSCO) version 4.0.5 (Simão et al., 2015) with a set of Actinopterygii orthologs (actinopterygii_odb10). The genes around the sex-associated region were predicted based on the protein sequences of model fish species in the Ensembl database (release 99) (Aken et al., 2017), namely *Danio rerio, Oryzias latipes, Gasterosteus aculeatus, Takifugu rubripes*, and *Tetraodon nigroviridis*, by Exonerate v2.4.0 (Slater and Birney, 2005). The PBT female genome sequence was obtained from the GenBank database (accession no. BKCK00000000). The nucleotide sequence comparison was performed by MUMmer v4 (Marcais et al., 2018). The repeat library of PBT genome was constructed by RepeatModeler Open-1.0, and the repetitive regions were masked by RepeatMasker Open-4.0 (http://www.repeatmasker.org). The duplicated regions were further detected by self-to-self BLASTN (Altschul et al., 1990) in each of the PBT male and female genomes, and the sequences detected were used also for masking.

### Genome-wide SNV/CNV analysis

The sequenced reads for 31 PBT individuals (15 males and 16 females) (Suda et al., 2019) were downloaded from the NCBI SRA database (accession no. DRR177387). The read sequences were extracted using SRA Toolkit (https://trace.ncbi.nlm.nih.gov/Traces/sra/sra.cgi?view=software), and then preprocessed using Trimmomatic v0.36 (Bolger et al., 2014) and ParDRe v2.2.5 (Gonzalez-Dominguez and Schmidt, 2016), according to Suda et al. Mapping to the scaffold sequences and single-nucleotide variant calling were conducted using MapCaller v0.9.9.37 (Lin and Hsu, 2019). After filtering out inadequate sites (not biallelic, depth < 20), genome-wide association (GWA) between males and females was estimated through 2 × 3 contingency tables using the R package, *rrBLUP* (Endelman, 2011), with minor allele frequency > 0.05. Statistical significance was defined as *P* < 0.05 after Bonferroni correction based on the total number of variant sites examined. For relative depth analysis, raw read depth per nucleotide site was counted using Samtools (*samtools depth*) (Li et al., 2009), and then divided by the median depth. Copy number variations (CNVs) were detected by CNVcaller (Wang et al., 2017), where the above-mentioned mapping data were scanned by a 500-bp window and regional copy numbers were obtained by merging them in the neighboring windows. The difference in the regional copy number between males and females was tested using the Mann-Whitney *U* test with Bonferroni correction. Furthermore, the regions carrying either of the three types of copy numbers (0, 0.5, or 1) among the 31 samples were selected, and the GWA was estimated in the same way as the SNV-based GWA scan mentioned above.

### Sequence analysis of sex-associated genes

The transcriptome data of southern bluefin tuna (Bar et al., 2016) were downloaded from the NCBI SRA database (accession no. SRP059929). The read sequences were extracted using SRA Toolkit, preprocessed by Trimmomatic, and assembled using Trinity v2.8.4 (Haas et al., 2013). From the contig sequences obtained, protein-coding sequences were predicted using TransDecoder v5.0.0 (Haas et al., 2013). Homologs of PBT genes were searched using BLASTP (Altschul et al., 1997) from the contig sequences, and the transcriptional variants were manually checked based on the sequence alignment using MAFFT v7.310 (Katoh et al., 2009). Using the Trinity contig sequences as a reference, gene expression was measured using RSEM v1.3.1 (Li and Dewey, 2011). For molecular phylogenetic analysis, homologs in other fish genomes were downloaded from the Ensembl database. The coding sequences were then aligned using MAFFT, where the deduced protein sequences were aligned and converted into nucleotide sequence alignments. The maximum-likelihood tree was constructed using RAxML-NG v0.9.0 (Kozlov et al., 2019).

## Results

### Genome assembly and validation of sex-associated markers

We constructed the genome sequence of male PBT by *de novo* assembly, yielding a total of 948 scaffolds (>1000 bp) and totaling 827 Mb (Table 1). Using BUSCO for quality assessment, this assembly completely captured 96.3% of Actinopterygii orthologs (3505 out of 3460). On the other hand, the score for the female genome was 89.3% (3253 out of 3640), consistent with previous reports. Using the sex-associated PCR primer sequences reported in a previous study, we performed a BLAST search against the current male genome, and found that two scaffolds, namely M175 and M44, carried the regions amplifiable by the primer sets (two male-specific sets, scaf64_3724604_F/R and scaf64_3726411_F/R, and a common primer set, scaf64_3724591_F/R) (Table 2). In scaffold M175 (232,647 bp in total), the regions amplified by the three primer sets were closely located. The estimated amplicon sizes were congruent with those previously observed only in males: 113 bp for scaf64_3724604_F/R, 143 bp for scaf64_3726411_F/R, and 142 bp for scaf64_3724591_F/R, respectively. Scaffold M44 (4,583,724 bp in total) had a single region amplified by scaf64_3724591_F/R, and the estimated amplicon size was 149 bp, which was equivalent to that observed previously in both males and females. Thus, based on the estimated PCR products, M175 carried all of the male-specific regions and M44 carried the common haplotype to both males and females. We compared the nucleotide sequences of M175 and M44, and found that M175 was almost dissimilar to that of M44 (Figure 1), while M44 was fully aligned with female scaffold F64 (Figure 2), in which the 6.5-kb sex-linked region (SLR) was reported in the previous study. The 6.5-kb SLR was found also in M44. On the other hand, we found no counterpart of M175 in the female genome, although the 6.5-kb SLR of M44/F64 (SLR_M44_ or SLR_F64_) was partially similar to the 10.7-kb region in M175 (SLR_M175_) (Figure 1). We assembled the male genome using Supernova2 and obtained scaffolds corresponding to M44 and M175, respectively (Supplementary Figure 1). The M44-like scaffold carrying SLR_M44_ was phased into two pseudo-haploid scaffolds, but SLR_M44_ itself was homozygotic. On the other hand, the scaffold carrying SLR_M175_ was not phased.

**Table 1.** Assembly statistics of PBT genomes.

|  | This study | Previous study* |
| --- | --- | --- |
| Number of scaffolds | 948 | 444 |
| Total scaffold size (Mbp) | 827 | 787 |
| N50 scaffold size (Mbp) | 13.2 | 7.9 |
| BUSCO completeness (%) | 96.3 | 89.3 |
| Single copy BUSCOs (%) | 93.6 | 86.1 |
| Duplicated BUSCOs (%) | 2.7 | 3.2 |

\*Accession no. BKCK00000000.

**Table 2.** Amplified sizes of sex identification PCR primer sets.

| Primer set | This study |  | Amplified size in the<br>previous study (bp) |  |
| --- | --- | --- | --- | --- |
|  | M44 | M175 | Female | Male |
| scaf64_3724604_F/R | - | 113 | - | 113 |
| scaf64_3726411_F/R | - | 143 | - | 143 |
| scaf64_3724591_F/R | 149 | 142 | 149 | 142, 149 |

**Figure 1.**
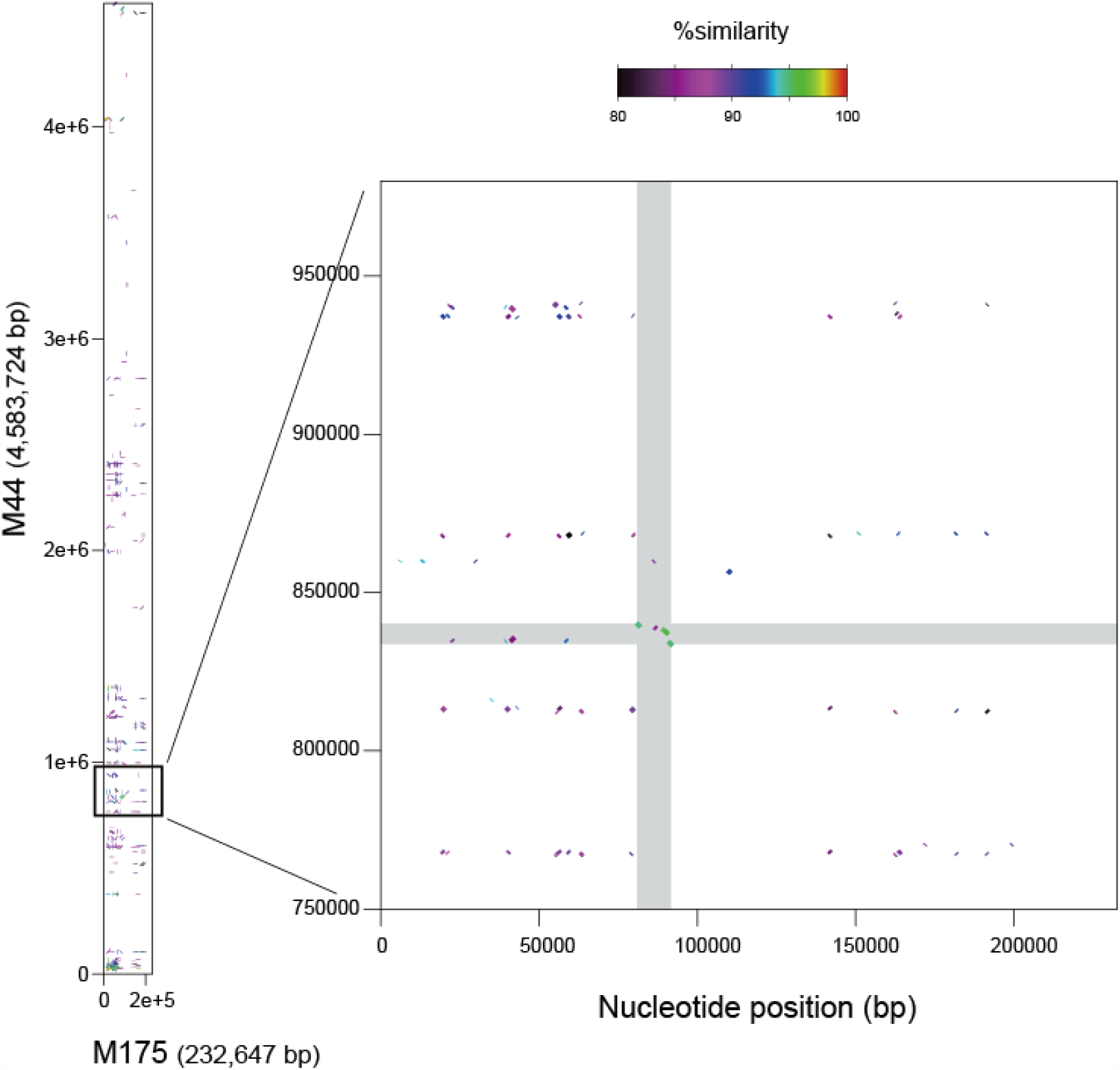
Sequence comparison between PBT male scaffolds M175 (X-axis) and M44 (Y-axis). Left panel indicates the full-size comparison (232,647 bp vs. 4,583,724 bp), and the boxed region is enlarged at right panel. In the right panel, SLR_M44_ and SLR_M175_ are highlighted in gray.

**Figure 2.**
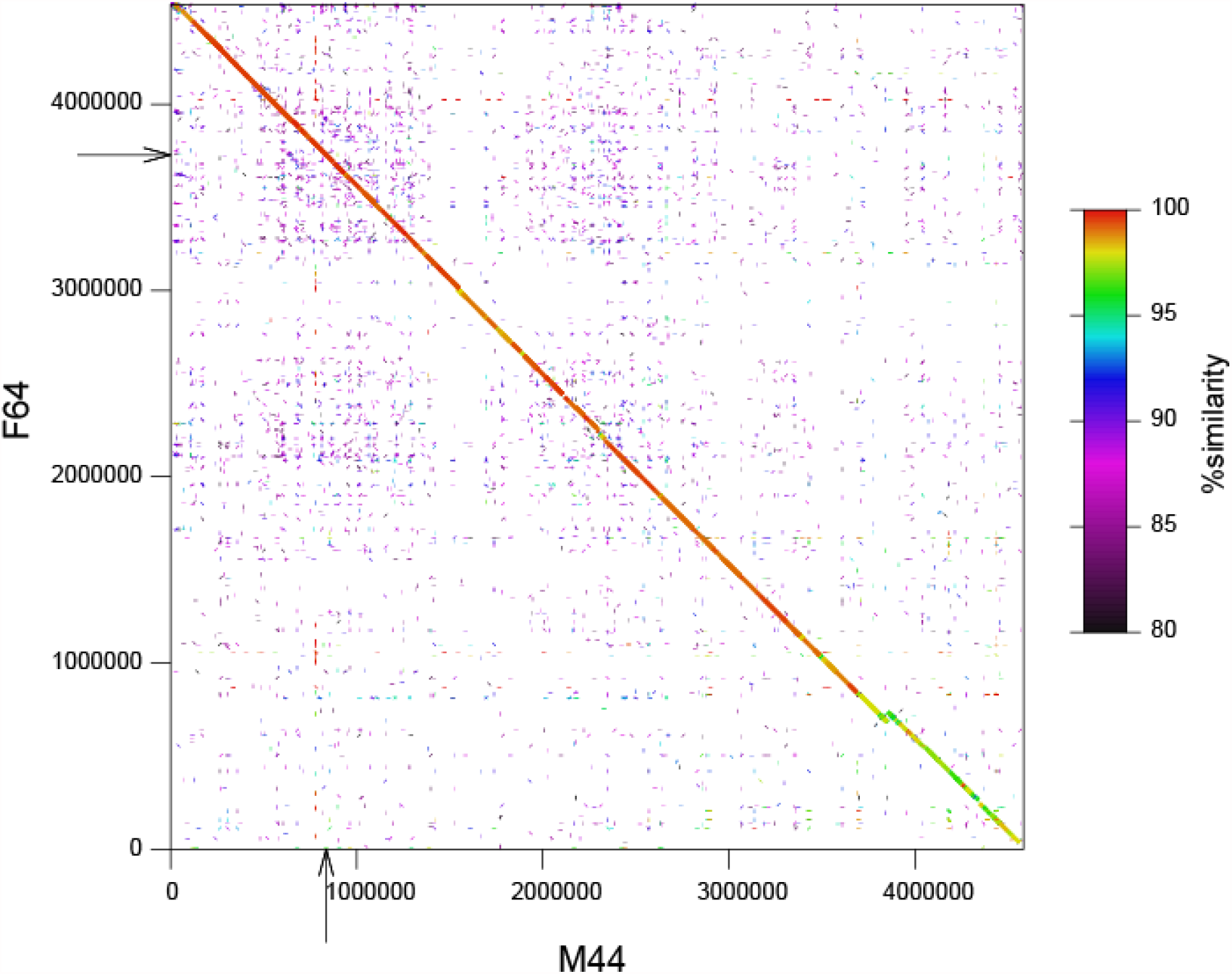
Sequence comparison between male scaffold M44 and female scaffold F64. The 6.5-kb SLRs are indicated with arrows, respectively.

### Read depth analysis

We downloaded the resequencing data of 31 PBT individuals (15 males and 16 females) and mapped the reads to the current male genome sequence. As a result, both male and female reads were fully mapped to SLR_M44_ (Figure 3 and Supplementary Figure 2). Regarding M175, the female reads were hardly mapped to SLR_M175_, in contrast to the male reads. Furthermore, there were consistently fewer male reads mapped to SLR_M175_ than SLR_M44_. We computed the median read depths for whole scaffolds, and thereby converted the read depths of SLR_M44_ and SLR_M175_ into the relative values. In males, the relative depths of SLR_M175_ were close to 0.5 (Figure 4). On the other hand, the relative depths of SLR_M44_ were close to 1 in both males and females. These observations strongly suggest that, in shotgun sequencing libraries, the DNA content derived from SLR_M175_ in the male samples is always half that derived from the average scaffolds, reflecting the ploidy or copy number. The majority of chromosomal regions, except for highly repetitive ones or those of polyploidized genomes, should exist as singletons in a diploid manner (2n), and the relative depths for many regions would be approximately 1. Therefore, our observations suggest that SLR_M175_, with a relative depth of ∼0.5, should exist in a haploid manner (1n) only in males, while SLR_M44_ should exist in a diploid manner, like many other regions, in both males and females.

**Figure 3.**
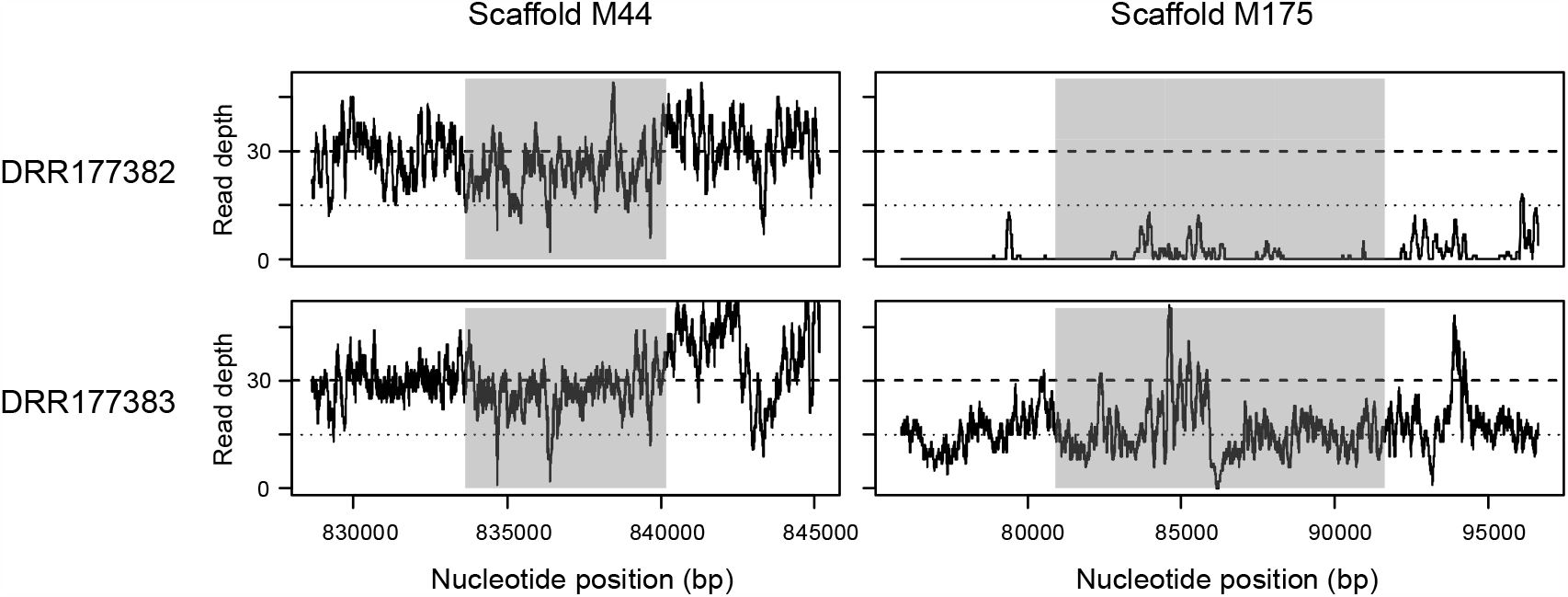
Mapped read depths around SLRs in resequenced PBT samples. For each of the female (DRR177382) and male (DRR177383) samples, SLR is highlighted in gray. The read depths for all the 31 samples are shown in Supplementary figure 2.

**Figure 4.**
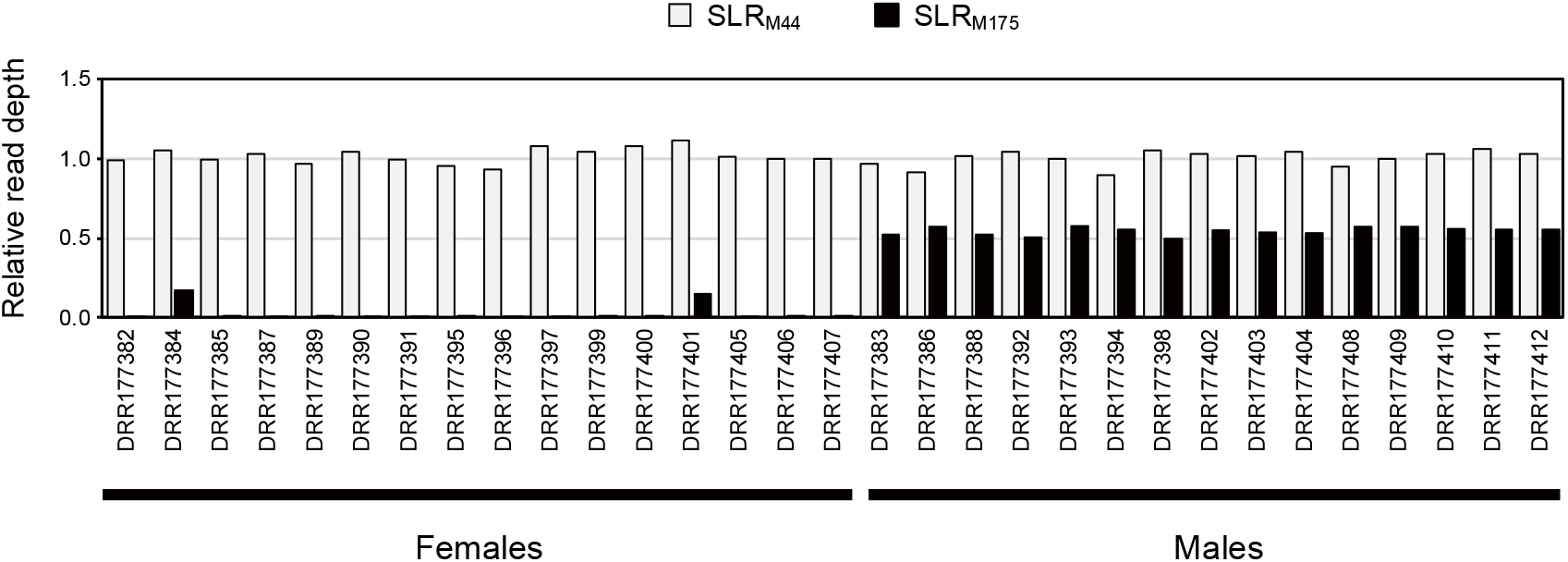
Comparison of relative read depths between males and females for SLR_M44_/SLR_M175_. The mapping of resequenced reads was conducted using an unmasked male reference.

### Genome-wide association analysis

Since the results obtained until now appeared to be inconsistent with those previously reported (see Discussion), we validated the previous mapping and GWA results based on the female genome. We mapped the 31 resequenced read sets to the female reference genome and re-performed GWA analysis. As a result, the previous result was roughly reproduced, with significant peaks observed in multiple scaffolds, including F64 (Figure 5A). However, when the mapped reads between the female and present male genomes were compared, over 10% of the reads mapped to female SLR_F64_ in the 15 male samples were found to be potentially derived from SLR_M175_ (Figure 6). This observation implies that the reads derived from the male-specific region, such as SLR_M175_, would have been incorrectly mapped to paralogous regions in the female genome in the previous study. To avoid the effect of such cross mapping, we performed the GWA scan again after the duplicated regions were thoroughly masked by RepeatMasker and BLASTN. This result was in contrast to that from the unmasked reference: no significant peaks were observed in the masked sequences (Figure 5B). Similarly, GWA scans were performed using the current male genome, but no significant peaks were found in the masked sequences (Supplementary Figure 3).

**Figure 5.**
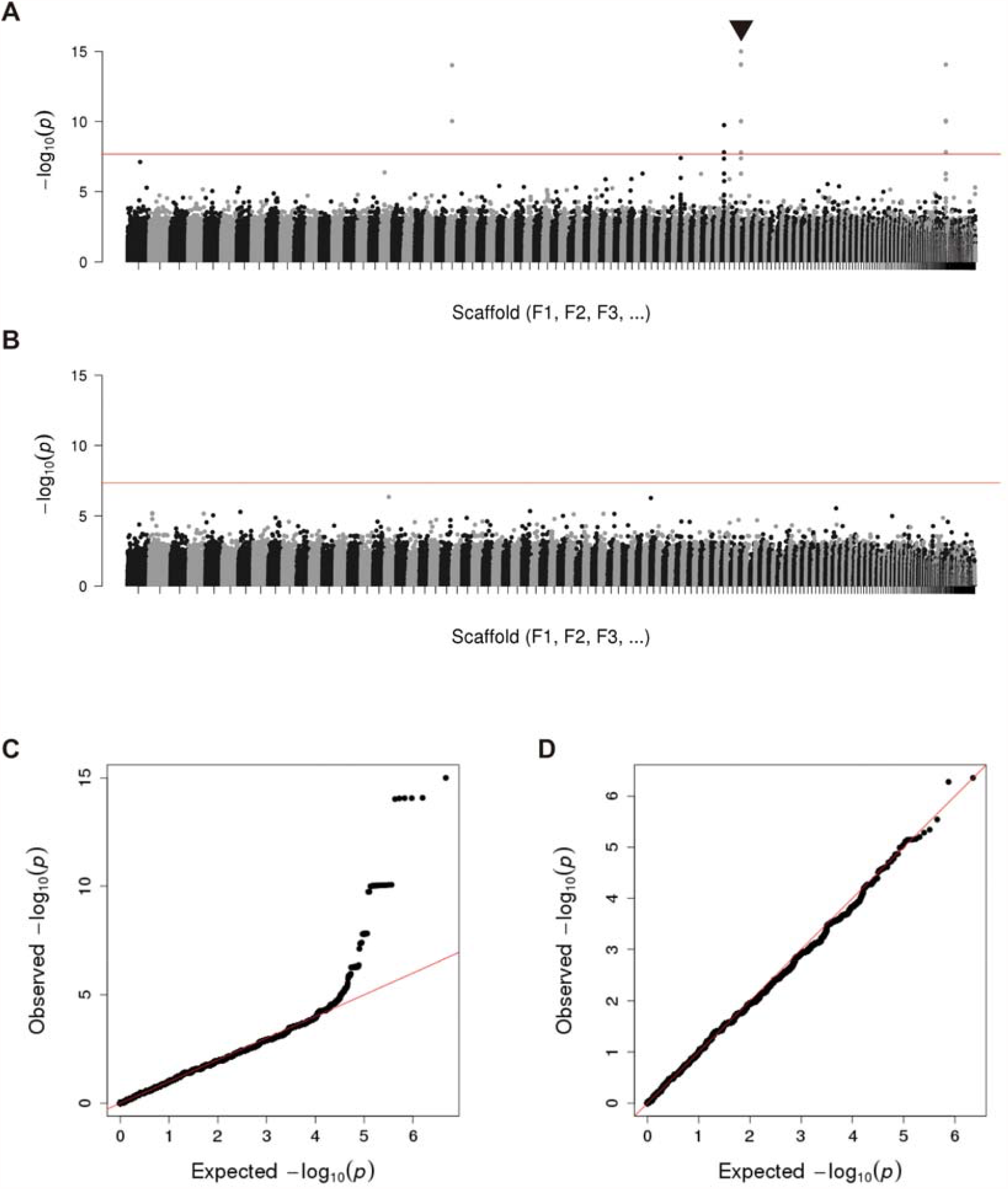
Genome-wide association with the sex of PBT using the female reference sequences. For the unmasked (A and C) and masked (B and D) references, Manhattan plot and Q-Q plot are shown, respectively. In Manhattan plots, genome-wide significance cutoffs based on Bonferroni correction are indicate by a red line, and the significant peak in scaffold F64 is indicated by a triangle. *P*-values lower than 10^−15^ are treated as 10^−15^ for convenience.

**Figure 6.**
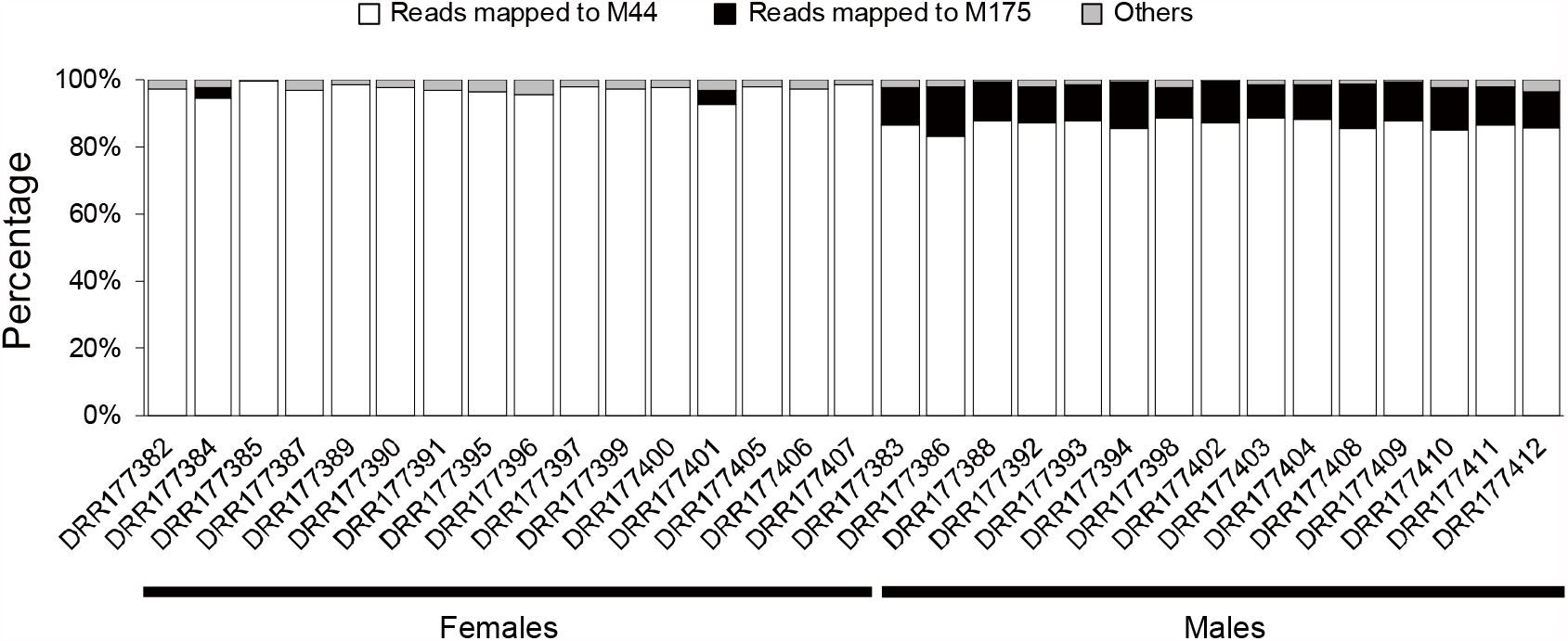
Classification of the resequenced reads mapped to SLR_F64_ in the female genome.

We then tested another approach to identify the regions carrying male-specific haplotypes, such as SLR_M175_. Using the mapping data for the 31 individuals, we computed copy number variations (CNVs) across the male genome and explored the regions whose copy numbers were significantly different between males and females. As a result, only one candidate was found in M175 (18,001–78,750 bp), very close to SLR_M175_, and the copy number in males was significantly larger than that in females (Figure 7A). All of the males had 0.5 copies of the detected region and all of the females had no copy (i.e., absent), in concordance with the relative depth analysis results (Figure 4). We computed genome-wide associations for the haploid regions by converting the copy numbers (0, 0.5, and 1) to three quasi-genotypes (0/0, 0/1, and 1/1). As a result, only one significant peak for this region was detected in scaffold M175 (Figure 7B).

**Figure 7.**
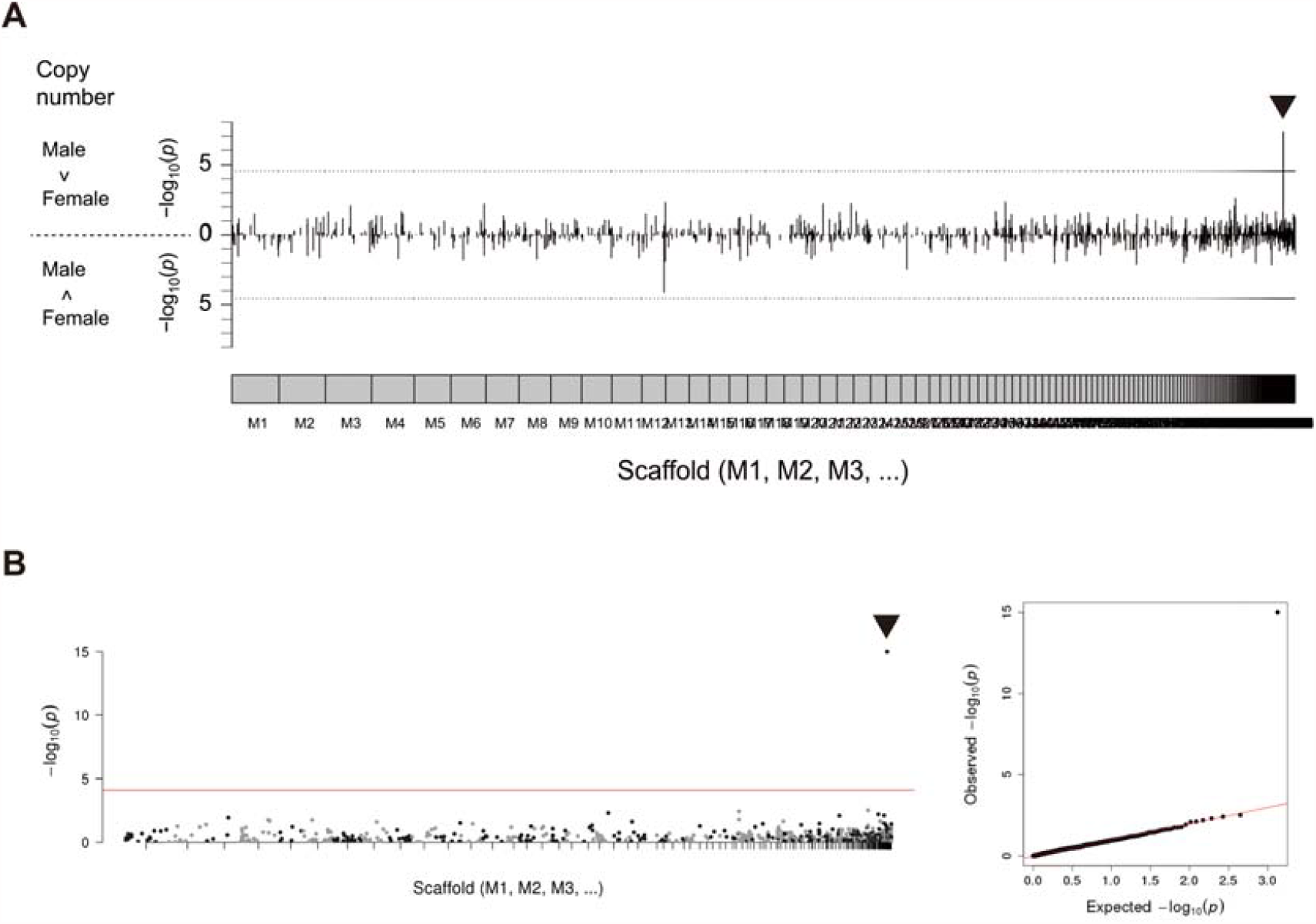
Genome-wide association with copy number between PBT males and females. The mapping of resequenced reads was conducted using a masked male reference. The plots on scaffold M175 are indicated by triangles. (A) Differences in regional copy number between males and females are denoted as the values of −log_10_(*p*) in the Mann-Whitney *U* test. Upward plots indicate that the copy number is larger in males than in females, while downward plots indicate the inverse. Genome-wide significance cutoffs based on Bonferroni correction are indicated by dashed lines. (B) Manhattan plot (left) and Q-Q plot (right) from CNV-based GWA scan. The regions with copy number 0, 0.5, or 1 were selected (*P*-values lower than 10^−15^ were treated as 10^−15^). In the Manhattan plot, genome-wide significance cutoff based on Bonferroni correction is indicated by a red line.

### Protein-coding genes in scaffold M175

We scanned the M175 sequence and predicted 12 protein-coding genes (Figure 8 and Supplementary Table 1). Of these, two genes, g3 (32,614–64,628 bp) and g4 (76,852–77,963 bp), were located in the haploid region (18,001–78,750 bp) detected by CNV analysis. The deduced amino acid sequence of g3 was similar to that of estrogen sulfotransferase encoded by *sult1st6* (e.g., with ∼83% identity to ENSORLP00000007539 of *Oryzias latipes*). Therefore, the homolog of PBT was named *sult1st6y*. In the case of g4, the deduced amino acid sequence was similar to that of ENSTRUP00000072273 (gene ID: ENSTRUG00000033234) of *Takifugu rubripes*, with ∼70% identity. As a “novel gene” in the Ensembl database, the function was unknown. However, the domain “Integrase, catalytic core” was predicted by Pfam and PROSITE, and found to overlap with the Gene3D domain “Ribonuclease H superfamily.” With regard to the neighboring genes, the protein sequence of g2 was matched to ENSTRUP00000068256 (ENSTRUG00000032453, novel gene), with the Pfam domain “PiggyBac transposable element-derived protein,” and g5 was matched to ENSTRUP00000067783 (ENSTRUG00000032600, novel gene) with the Pfam domain “Reverse transcriptase/retrotransposon-derived protein, RNase H-like domain.” Notably, g1 was similar to g5 and matched to ENSTRUP00000067783, suggesting that g2–g4 were sandwiched between duplicated genes (g1 and g5). In addition, g6 was matched to ENSORLP00000040498 (ENSORLG00000025541, novel gene) with the Pfam domain “Transposase, Tc1-like,” and the other genes, g7–g12, were matched to the same protein, ENSORLP00000044417 (ENSORLG00000028531, novel gene) with the Gene3D domain “Immunoglobulin-like fold.” Thus, only g3 was a functionally annotated gene, and the others were novel or related to transposable elements. We found no homologs of *sult1st6y* (g3) in the scaffolds M44/F64. Instead, the paralog was found in male scaffold 30 (M30, 6594208–6602041 bp), named *sult1st6a*, and the nucleotide and protein identities to *sult1st6y* were 92% and 89%, respectively. Male scaffold M30 corresponded to a part of female scaffold 47 (F47) (Supplementary Figure 4), where *sult1st6a* was also found.

**Figure 8.**
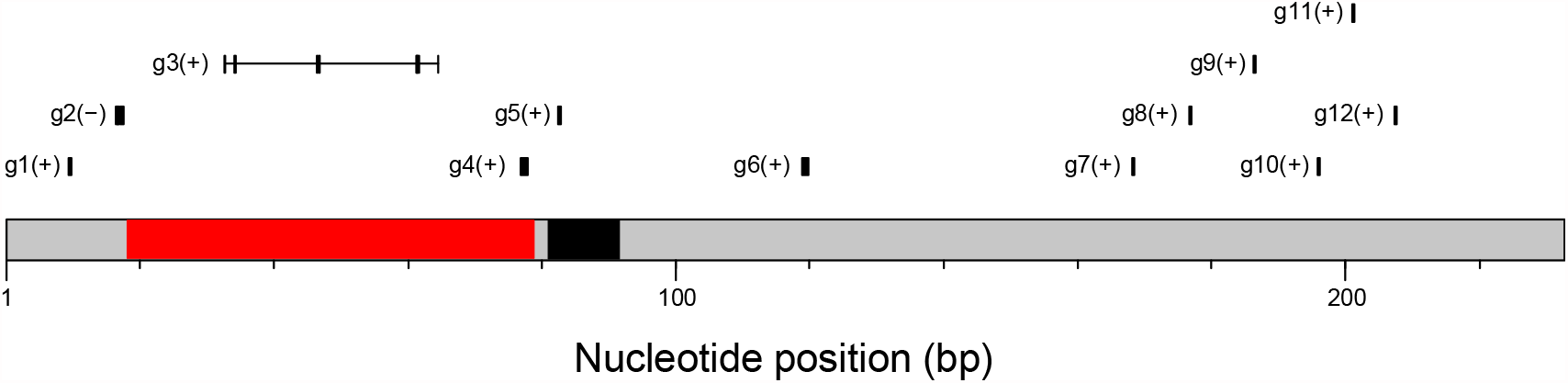
Protein-coding regions in scaffold M175. The region predicted with a copy number of 0.5 and 10.7-kb SLR_M175_ are shown in red and black, respectively.

To determine whether these *sult1st6* genes are present in other tuna species, we downloaded the RNA-seq data of southern bluefin tuna (SBT), which were obtained from 10 male testis samples and 10 female ovary samples. We assembled the reads using Trinity and conducted a homology search against the contigs using PBT *sult1st6y*/*sult1st6a* sequences. As a result, we found the homologs of *sult1st6y*/*sult1st6a* in SBT, although the Trinity contigs were originally classified into a single cluster. We conducted phylogenetic analysis using PBT and SBT *sult1st6* and the homologs in the Ensembl database (Figure 9 and Supplementary Figure 5). The phylogenetic tree showed that each SBT homolog formed a clade with each of PBT’s *sult1st6y*/*sult1st6a*, respectively, suggesting that SBT also had two paralogs of *sult1st6* and the duplication to *sult1st6y* and *sult1st6a* occurred in the common ancestor of PBT and SBT. Using the transcript contigs as a reference, we mapped the reads in each of the 20 samples to those and measured the gene expression levels (Figure 10). As a whole, the expression of SBT’s *sult1st6y*/*sult1st6a* was low compared to that of the control (0–2% of β-actin in FPKM). In detail, the expression of *sult1st6a* was observed in both the testis and ovary samples, although it tended to be low in the latter (0–0.01% of β-actin in FPKM) (Figure 10A). The expression of *sult1st6y* was observed in all 10 testis samples, but none of ovary samples, indicating a significant bias of *sult1st6y* expression between the testes and ovaries (*P* < 0.05, Fisher’s exact test) (Figure 10B).

**Figure 9.**
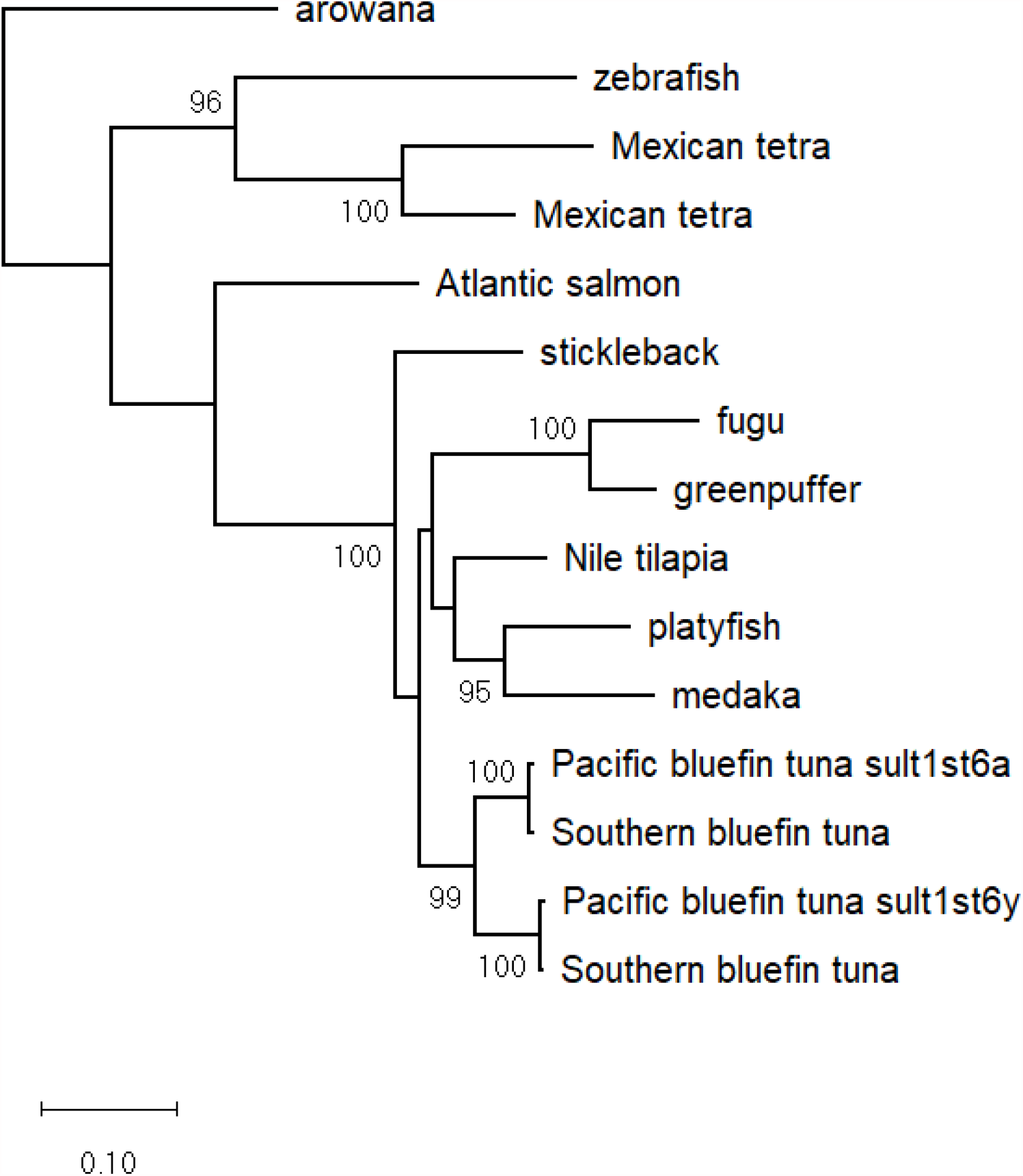
Phylogenetic tree of *sult1st6* in fish species. For each of the nodes, the bootstrap probability is indicated when it is >95%. The tree constructed using more homologs linked with the Ensembl IDs is shown in Supplementary Figure 5.

**Figure 10.**
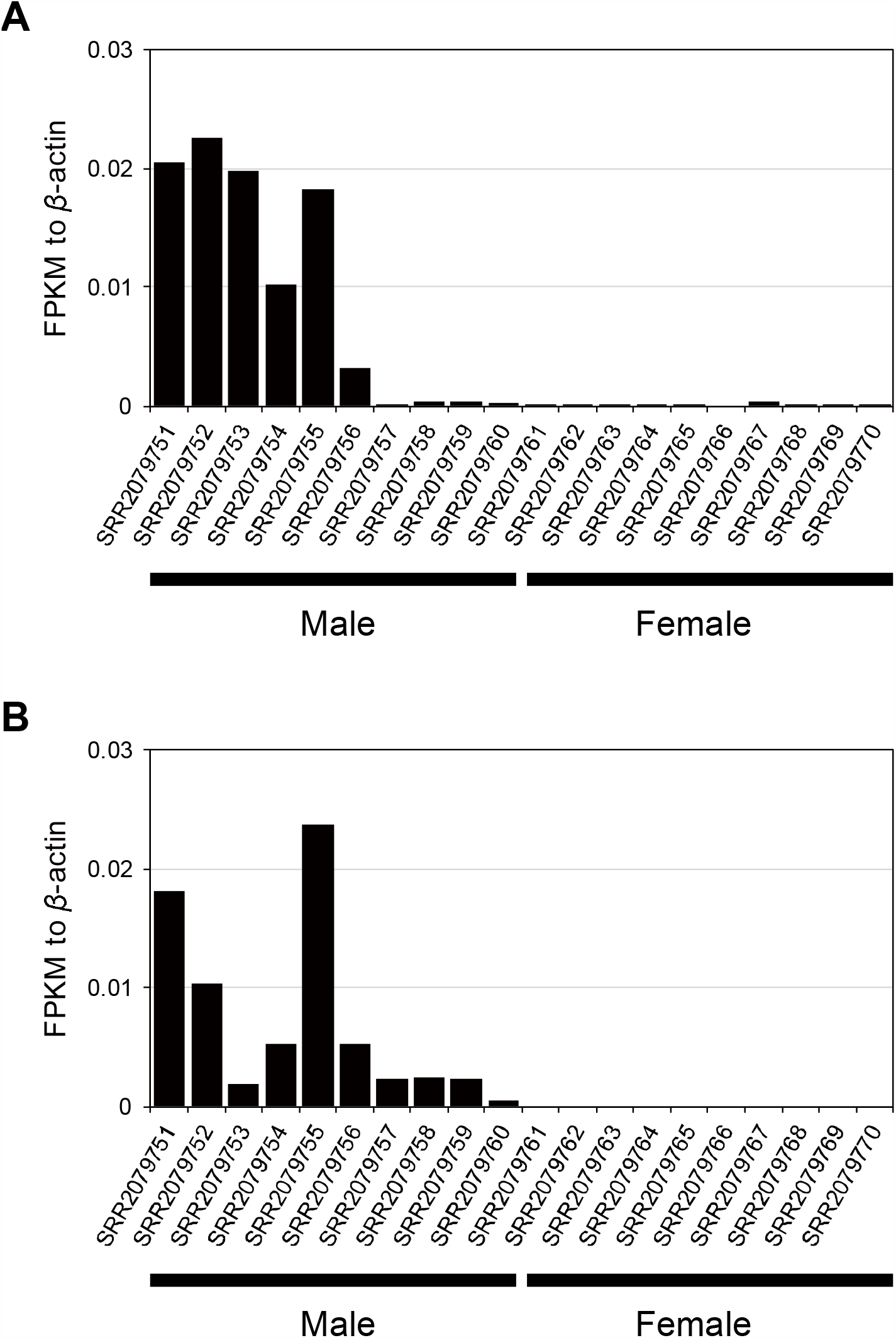
Expression of *sult1st6* in southern bluefin tuna’s gonad cells. (A) *sult1st6a* and (B) *sult1st6y*.

## Discussion

In this study, the male genome of Pacific bluefin tuna was sequenced using long-read sequencing technologies. The scaffold sequences assembled were more contiguous than those published in 2013, which were obtained from pyrosequencing and short paired-end reads. Completeness assessment indicates that the current version is better than that of the recent female genome, although the number of scaffolds (948) is larger than that for the female genome (444). Therefore, the current genome is expected to be useful for the validation and extension of previous results regarding the sex identification of PBT. In the previous study, three PCR primer sets were designed from the sequence of female scaffold F64, so that the amplification patterns were different between PBT males and females. In this study, we first searched for the locus amplified by the primer sets in the current male genome, and found two regions: one in scaffold M44 and the other in M175. Based on the results of the large-scale sequence comparison, M44 was found to be the counterpart of F64, while M175 was male-specific. Since M44 and F64 were well aligned with each other, including 6.5-kb SLRs (SLR_M44_/SLR_F64_), it is unlikely that large-scale misassemblies are present in M44/F64. The PCR amplification pattern in SLR_M44_, estimated using primer sequences, was identical to that observed in females. Regarding M175, all the PCR amplicons with male-specific sizes were observed within the 10.7-kb region (SLR_M175_). It is worth noting that the PCR primer sets were originally designed based on the SNVs predicted in F64 by GWA analysis. Therefore, according to the previous results, M44 and M175 should be allelic to each other. However, these sequences are quite dissimilar to each other. Furthermore, based on the mapping result of resequenced data, the relative read depths to that for the average scaffold data were (SLR_M44_, SLR_M175_) ≈ (1, 0) in females but (SLR_M44_, SLR_M175_) ≈ (1, 0.5) in males. These estimates indicate that SLR_M44_ should exist in a diploid manner in both males and females, whereas SLR_M175_ should exist in a haploid manner only in males. If SLR_M44_ was allelic to SLR_M175_ in males, the relative read depths (SLR_M44_, SLR_M175_) should be (0.5, 0.5). The result of relative read depth is congruent with that of phased assembly by Supernova2, where two pseudo-haplotype scaffolds carrying homozygotic SLR_M44_ were obtained, but only unphased one was obtained for SLR_M175_. These results indicate that SLR_M44_ and SLR_M175_ are located at different loci in the male genome, where SLR_M175_ is the male-specific structural variant (i.e., inserted in males or deleted in females) and is inherited paternally. In other words, SLR_F64_ (or SLR_M44_) reported in the previous study was not associated with PBT sex, although the PCR primer sets designed from SLR_F64_ are still available for sex identification based on the amplification pattern in SLR_M175_.

Now, the previous result, in which male-specific heterozygous polymorphisms were detected in F64, needs to be reconsidered. The previous GWA scan was performed using the female genome as a reference, focusing on male-heterozygous SNVs or small indels, which may be detectable by the resequenced short reads. However, M175 found in the current study is a large-scale structural variant absent from the female genome and should be undetectable by short-read mapping. We considered the possibility that the reads derived from paralogous loci were cross-mapped to the female reference genome, causing fake heterozygous SNV calls in genotyping prior to GWA analysis. In the previous study, the repeat or paralogous sequences were not masked; therefore, it is likely that the reads derived from the paternal 10.7-kb region (i.e., SLR_M175_) were mapped to the 6.5-kb region of F64 in the analysis of male DNA samples. This hypothesis was confirmed by checking the mapped reads: at least 10% of the male reads mapped to SLR_F64_ in the previous analysis were estimated to be derived from SLR_M175_. Furthermore, we compared the GWA results between the two conditions for the female reference sequence: duplicated regions were unmasked (i.e., raw sequences) and masked. The result was clear-cut, where the peaks in multiple scaffolds were roughly reproduced in the unmasked sequences, but no such peaks were observed in the masked sequences. Thus, we conclude that the previous GWA result was an artifact due to cross-mapping, and many of the scaffolds predicted (e.g., F64) were not associated with the sex of PBT. Since we found no significant GWA peaks in the masked male reference genome, the SLR of PBT would be undetectable from the information of SNVs or small indels. The current results provide a caveat that traditional GWA analysis, which targets SNVs using short reads, may not always be sufficient for the identification of SLR in species with a genetic sex determination system.

Instead of SLR_M44_, we found that SLR_M175_ is a male-specific locus absent from the female genome. It should be noted that this finding is due to a homology search of the primer sequences designed in the previous study. Therefore, it is possible that the current male genome contains other scaffolds carrying sex-associated structural variants. To check this possibility, we conducted a genome-wide scan based on a CNV detection scheme using mapping data. As a result, we observed that only M175 had a 61-kb region (18,001–78,750 bp) where the CNV was significantly different between males and females. Since this region is very close to SLR_M175_, eventually the candidate sex-associated locus was narrowed down to a single region around or including SLR_M175_. Regarding the CNV detected, we found that PBT females have no copy, while males have 0.5 copies of the region, indicating that this region exists in a haploid manner in males, or equivalently, that this region is paternally inherited. Thus, we conclude that this region (61-kb region + SLR_M175_) is the only candidate for the Y-linked locus in PBT. In M175, we predicted a total of 12 protein-coding genes, two of which (g3 and g4) were included in the predicted haploid region, one (g3) of which was annotated as encoding estrogen sulfotransferase (SULT1ST6), while the other (g4) was functionally unknown. We note that g4 has the domain “Integrase, catalytic core” or “Ribonuclease H superfamily” which is a feature of the retroviral integrase superfamily (Nowotny, 2009). Regarding the other 10 predicted genes in M175, the functions are unknown. In particular, g1, g2, and g5 may be involved in transposons according to domain prediction. Therefore, around the male-specific haploid region, only the centermost g3, namely *sult1st6y*, is functionally annotated and sandwiched between possibly transposable elements, implying that the g3 locus may have been unstable and transposed from the ancestral locus.

The current study suggests that *sult1st6y* is the most convincing gene involved in the sex determination or differentiation of PBT. The encoding protein, SULT1ST6, belongs to the family of cytosolic sulfotransferases (SULTs) and has been studied as a detoxifying enzyme in mammals (Suiko et al., 2017). Regarding fish species, the SULT genes have been investigated mainly in zebrafish, and 20 distinct paralogs (9 SULT1s, 3 SULT2s, 5 SULT3s, 1 SULT4, 1 SULT5, and 1 SULT6) have been characterized. Among these, SULT1ST6, one of the nine SULT1 paralogs, displays strict substrate specificity for estrogens (e.g., estrone). This gene is thought to be involved in organogenesis, such as the development of the eye and muscle in zebrafish (Yasuda et al., 2005). On the other hand, there is no report about the relationship between estrogen sulfotransferase and sex development. In fish, steroid hormones are considered to be the key factor for sex determination (Yamamoto, 1969; Devlin and Nagahama, 2002; Nakamura, 2010), wherein sex can change depending on the levels of estrogen or androgen production in gonad cells. The gene directly contributing to steroidogenesis has also been reported as a sex-determination gene in yellowtails (Koyama et al., 2019). Since endogenous estrogens are inactivated by sulfation (Raftogianis et al., 2000), it may not be surprising that the function of *sult1st6* is involved in sex development. If this gene is expressed in gonad cells at the initiation of sex development, it may inhibit female development by depleting active estrogens, resulting in male development. In the present study, we found that *sult1st6y* of PBT was paternally inherited in the population. It is unlikely that an estrogen-inactivating gene with such a mode of inheritance is unrelated to the sex development of PBT. In addition, we found that both female and male PBT had a paralog of *sult1st6y*, namely *sult1st6a*, in male scaffold M30 or female scaffold F47. It is worth noting that, in the previous study, F47 was another candidate carrying sex-associated SNVs (Suda et al., 2019), and the locus of *sult1st6a* was close to the artifact GWA peak. This observation is not only attributable to cross-mapping, as mentioned before, but also indirectly suggests that the *sult1st6*-like gene might be associated with the sex of PBT. Since *sult1st6a* is inherited regardless of sex in a diploid manner, we speculate that *sult1st6a* plays a common role in PBT males and females, for example, in organogenesis at the early developmental stage, as proposed in the model fish. This suggests that *sult1st6y* could have been immune from the original role of *sult1st6* as a redundant copy: after gene duplication in the ancestor of PBT, the function of *sult1st6y* may have been spatiotemporally differentiated from that of *sult1st6a* in the proto-paternal line. In fact, it has been reported that a copy of duplicated genes may be sex-determination gene in fishes (Hattori et al., 2012; Yano et al., 2012; Matsuda and Sakaizumi, 2016). We found that homologs of *sult1st6y* and *sult1st6a* were present in the southern bluefin tuna. Phylogenetic analysis suggested that gene duplication occurred in the common ancestor of PBT and SBT. The duplication event is specific to the tuna lineage, not observed in other fishes, except for tandem duplications in some species. According to the transcriptome, SBT *sult1st6y* was expressed in male’s testis but not in female’s ovary. Considering that the expression of *sult1st6a* was at least detectable in females, although at a much lower level than in males, *sult1st6y* below the detection limit might be originally absent from SBT females like PBT females. Although this hypothesis will need to be tested by sequencing the entire SBT genome, our observations suggest that SBT *sult1st6y* may also be involved in sex development like PBT *sult1st6y*.

The results presented in this study provide an attractive hypothesis for *sult1st6y* being a sex-determination gene in *Thunnus* fishes, or at least functioning at an important point in the sex-differentiation cascade. Our hypothesis may be tested by further genetic experiments, although the handling of tuna in tanks is not easy. Recently, genome-editing technology has been applied to the mutagenesis of tuna (Higuchi et al., 2019). For example, it may be possible to check whether the sex ratio of larvae is heavily biased by the knockout of *sult1st6y* in fertilized eggs. Lastly, it is worth emphasizing that the CNV-based approach used in this study is very useful for identifying a trait-associated region using short reads when it is involved in large structural variants. In particular, when the region contains recently duplicated sequences, the traditional SNV-based approach may not only miss structural variants but also produce artifacts due to cross-mapping to unassociated regions. In other fish species, sex-associated SNVs have often been observed in multiple scaffolds (Fowler and Buonaccorsi, 2016; Star et al., 2016; Dong et al., 2019), some of which might be cases of cross-mapping of short reads, unless the sexes are polygenically determined. We believe that a combination of traditional SNV- and CNV-based approaches is useful for the identification of sex-associated genes in fish species with a genetic sex-determination system.

## Data availability statement

The genome sequences of Pacific bluefin tuna obtained in this study were deposited to the DNA Data Bank of Japan. The accession numbers can be found in the article.

## Author contributions

YN and AF conceived the study. KG coordinated the experiments for sampling, and KH, KK, MY, and TT performed those. AF conducted genome sequencing, and YN conducted bioinformatic analyses. YN drafted the manuscript, and MY helped to improve it. All authors read and approved the final version.

## Acknowledgements

This work was supported by the Japan Fisheries Research and Education Agency (No. 3BA101). We would like to thank Editage (www.editage.jp) for English language editing.

## References

Agawa, Y., Iwaki, M., Komiya, T., Honryo, T., Tamura, K., Okada, T., et al. (2015). Identification of male sex-linked DNA sequence of the cultured Pacific bluefin tuna Thunnus orientalis. Fisheries Science 81(1), 113–121. doi: 10.1007/s12562-014-0833-8.

Aken, B.L., Achuthan, P., Akanni, W., Amode, M.R., Bernsdorff, F., Bhai, J., et al. (2017). Ensembl 2017. Nucleic Acids Res 45(D1), D635–D642. doi: 10.1093/nar/gkw1104.

Altschul, S.F., Gish, W., Miller, W., Myers, E.W., and Lipman, D.J. (1990). Basic local alignment search tool. J Mol Biol 215(3), 403–410. doi: 10.1016/S0022-2836(05)80360-2S0022-2836(05)80360-2[pii].

Altschul, S.F., Madden, T.L., Schaffer, A.A., Zhang, J., Zhang, Z., Miller, W., et al. (1997). Gapped BLAST and PSI-BLAST: a new generation of protein database search programs. Nucleic Acids Res 25(17), 3389–3402. doi: gka562[pii].

Bar, I., Cummins, S., and Elizur, A. (2016). Transcriptome analysis reveals differentially expressed genes associated with germ cell and gonad development in the Southern bluefin tuna (Thunnus maccoyii). BMC Genomics 17, 217. doi: 10.1186/s12864-016-2397-8.

Bolger, A.M., Lohse, M., and Usadel, B. (2014). Trimmomatic: a flexible trimmer for Illumina sequencedata.Bioinformatics30(15),2114–2120.doi: 10.1093/bioinformatics/btu170.

Chin, C.S., Alexander, D.H., Marks, P., Klammer, A.A., Drake, J., Heiner, C., et al. (2013). Nonhybrid, finished microbial genome assemblies from long-read SMRT sequencing data. Nat Methods 10(6), 563–569. doi: 10.1038/nmeth.2474.

Coombe, L., Zhang, J., Vandervalk, B.P., Chu, J., Jackman, S.D., Birol, I., et al. (2018). ARKS: chromosome-scale scaffolding of human genome drafts with linked read kmers. BMC Bioinformatics 19(1), 234. doi: 10.1186/s12859-018-2243-x.

Cui, Z., Liu, Y., Wang, W., Wang, Q., Zhang, N., Lin, F., et al. (2017). Genome editing reveals dmrt1 as an essential male sex-determining gene in Chinese tongue sole (Cynoglossus semilaevis). Sci Rep 7, 42213. doi: 10.1038/srep42213.

Devlin, R.H., and Nagahama, Y. (2002). Sex determination and sex differentiation in fish: an overview of genetic, physiological, and environmental influences. Aquaculture 208(3), 191–364. doi: https://doi.org/10.1016/S0044-8486(02)00057-1.

Dong, C., Jiang, P., Zhang, J., Li, X., Li, S., Bai, J., et al. (2019). High-Density Linkage Map and Mapping for Sex and Growth-Related Traits of Largemouth Bass (Micropterus salmoides). Front Genet 10, 960. doi: 10.3389/fgene.2019.00960.

Endelman, J.B. (2011). Ridge Regression and Other Kernels for Genomic Selection with R Package rrBLUP. The Plant Genome 4(3), 250–255. doi: 10.3835/plantgenome2011.08.0024.

Fowler, B.L., and Buonaccorsi, V.P. (2016). Genomic characterization of sex-identification markers in Sebastes carnatus and Sebastes chrysomelas rockfishes. Mol Ecol 25(10), 2165–2175. doi: 10.1111/mec.13594.

Gonzalez-Dominguez, J., and Schmidt, B. (2016). ParDRe: faster parallel duplicated reads removal tool for sequencing studies. Bioinformatics 32(10), 1562–1564. doi: 10.1093/bioinformatics/btw038.

Haas, B.J., Papanicolaou, A., Yassour, M., Grabherr, M., Blood, P.D., Bowden, J., et al. (2013). De novo transcript sequence reconstruction from RNA-seq using the Trinity platform for reference generation and analysis. Nat Protoc 8(8), 1494–1512. doi: 10.1038/nprot.2013.084.

Hattori, R.S., Murai, Y., Oura, M., Masuda, S., Majhi, S.K., Sakamoto, T., et al. (2012). A Y-linked anti-Mullerian hormone duplication takes over a critical role in sex determination. Proc Natl Acad Sci U S A 109(8), 2955–2959. doi: 10.1073/pnas.1018392109.

Higuchi, K., Kazeto, Y., Ozaki, Y., Yamaguchi, T., Shimada, Y., Ina, Y., et al. (2019). Targeted mutagenesis of the ryanodine receptor by Platinum TALENs causes slow swimming behaviour in Pacific bluefin tuna (Thunnus orientalis). Sci Rep 9(1), 13871. doi: 10.1038/s41598-019-50418-3.

Jackman, S.D., Coombe, L., Chu, J., Warren, R.L., Vandervalk, B.P., Yeo, S., et al. (2018). Tigmint: correcting assembly errors using linked reads from large molecules. BMC Bioinformatics 19(1), 393. doi: 10.1186/s12859-018-2425-6.

Kamiya, T., Kai, W., Tasumi, S., Oka, A., Matsunaga, T., Mizuno, N., et al. (2012). A trans-species missense SNP in Amhr2 is associated with sex determination in the tiger pufferfish, Takifugu rubripes (fugu). PLoS Genet 8(7), e1002798. doi: 10.1371/journal.pgen.1002798.

Katoh, K., Asimenos, G., and Toh, H. (2009). Multiple alignment of DNA sequences with MAFFT. Methods Mol Biol 537, 39–64. doi: 10.1007/978-1-59745-251-9_3.

Koyama, T., Nakamoto, M., Morishima, K., Yamashita, R., Yamashita, T., Sasaki, K., et al. (2019). A SNP in a steroidogenic enzyme is associated with phenotypic sex in Seriola fishes. Curr Biol 29(11), 1901–1909 e1908. doi: 10.1016/j.cub.2019.04.069.

Kozlov, A.M., Darriba, D., Flouri, T., Morel, B., and Stamatakis, A. (2019). RAxML-NG: a fast, scalable and user-friendly tool for maximum likelihood phylogenetic inference. Bioinformatics 35(21), 4453–4455. doi: 10.1093/bioinformatics/btz305.

Li, B., and Dewey, C.N. (2011). RSEM: accurate transcript quantification from RNA-Seq data with or without a reference genome. BMC Bioinformatics 12, 323. doi: 10.1186/1471-2105-12-323.

Li, H., Handsaker, B., Wysoker, A., Fennell, T., Ruan, J., Homer, N., et al. (2009). The Sequence Alignment/Map format and SAMtools. Bioinformatics 25(16), 2078–2079. doi: 10.1093/bioinformatics/btp352.

Lin, H.-N., and Hsu, W.-L. (2019). MapCaller – An integrated and efficient tool for short-read mapping and variant calling using high-throughput sequenced data. bioRxiv, 783605. doi: 10.1101/783605.

Marcais, G., Delcher, A.L., Phillippy, A.M., Coston, R., Salzberg, S.L., and Zimin, A. (2018). MUMmer4: A fast and versatile genome alignment system. PLoS Comput Biol 14(1), e1005944. doi: 10.1371/journal.pcbi.1005944.

Matsuda, M., Nagahama, Y., Shinomiya, A., Sato, T., Matsuda, C., Kobayashi, T., et al. (2002). DMY is a Y-specific DM-domain gene required for male development in the medaka fish. Nature 417(6888), 559–563. doi: 10.1038/nature751.

Matsuda, M., and Sakaizumi, M. (2016). Evolution of the sex-determining gene in the teleostean genus Oryzias. Gen Comp Endocrinol 239, 80–88. doi: 10.1016/j.ygcen.2015.10.004.

Nakamura, M. (2010). The mechanism of sex determination in vertebrates-are sex steroids the key-factor? J Exp Zool A Ecol Genet Physiol 313(7), 381–398. doi: 10.1002/jez.616.

Nakamura, Y., Mori, K., Saitoh, K., Oshima, K., Mekuchi, M., Sugaya, T., et al. (2013). Evolutionary changes of multiple visual pigment genes in the complete genome of Pacific bluefin tuna. Proc Natl Acad Sci U S A 110(27), 11061–11066. doi: 10.1073/pnas.1302051110.

Nanda, I., Kondo, M., Hornung, U., Asakawa, S., Winkler, C., Shimizu, A., et al. (2002). A duplicated copy of DMRT1 in the sex-determining region of the Y chromosome of the medaka, Oryzias latipes. Proc Natl Acad Sci U S A 99(18), 11778–11783. doi: 10.1073/pnas.182314699.

Nowotny, M. (2009). Retroviral integrase superfamily: the structural perspective. EMBO Rep 10(2), 144–151. doi: 10.1038/embor.2008.256.

Pan, Q., Feron, R., Yano, A., Guyomard, R., Jouanno, E., Vigouroux, E., et al. (2019). Identification of the master sex determining gene in Northern pike (Esox lucius) reveals restricted sex chromosome differentiation. PLoS Genet 15(8), e1008013. doi: 10.1371/journal.pgen.1008013.

Pandian, T.J. (2011). Sex Determination in Fish. CRC Press.

Pennell, M.W., Mank, J.E., and Peichel, C.L. (2018). Transitions in sex determination and sex chromosomes across vertebrate species. Mol Ecol 27(19), 3950–3963. doi: 10.1111/mec.14540.

Raftogianis, R., Creveling, C., Weinshilboum, R., and Weisz, J. (2000). Estrogen metabolism by conjugation. J Natl Cancer Inst Monogr (27), 113–124. doi: 10.1093/oxfordjournals.jncimonographs.a024234.

Simão, F.A., Waterhouse, R.M., Ioannidis, P., Kriventseva, E.V., and Zdobnov, E.M. (2015). BUSCO: assessing genome assembly and annotation completeness with single-copy orthologs. Bioinformatics 31(19), 3210–3212. doi: 10.1093/bioinformatics/btv351.

Slater, G.S., and Birney, E. (2005). Automated generation of heuristics for biological sequence comparison. BMC Bioinformatics 6, 31. doi: 1471-2105-6-31[pii] 10.1186/1471-2105-6-31.

Star, B., Torresen, O.K., Nederbragt, A.J., Jakobsen, K.S., Pampoulie, C., and Jentoft, S. (2016). Genomic characterization of the Atlantic cod sex-locus. Sci Rep 6, 31235. doi: 10.1038/srep31235.

Suda, A., Nishiki, I., Iwasaki, Y., Matsuura, A., Akita, T., Suzuki, N., et al. (2019). Improvement of the Pacific bluefin tuna (Thunnus orientalis) reference genome and development of male-specific DNA markers. Sci Rep 9(1), 14450. doi: 10.1038/s41598-019-50978-4.

Suiko, M., Kurogi, K., Hashiguchi, T., Sakakibara, Y., and Liu, M.C. (2017). Updated perspectives on the cytosolic sulfotransferases (SULTs) and SULT-mediated sulfation. Biosci Biotechnol Biochem 81(1), 63–72. doi: 10.1080/09168451.2016.1222266.

Walker, B.J., Abeel, T., Shea, T., Priest, M., Abouelliel, A., Sakthikumar, S., et al. (2014). Pilon: an integrated tool for comprehensive microbial variant detection and genome assembly improvement. PLoS One 9(11), e112963. doi: 10.1371/journal.pone.0112963.

Wang, X., Zheng, Z., Cai, Y., Chen, T., Li, C., Fu, W., et al. (2017). CNVcaller: highly efficient and widely applicable software for detecting copy number variations in large populations. Gigascience 6(12), 1–12. doi: 10.1093/gigascience/gix115.

Warren, R.L., Yang, C., Vandervalk, B.P., Behsaz, B., Lagman, A., Jones, S.J., et al. (2015). LINKS: Scalable, alignment-free scaffolding of draft genomes with long reads. Gigascience 4, 35. doi: 10.1186/s13742-015-0076-3.

Weisenfeld, N.I., Kumar, V., Shah, P., Church, D.M., and Jaffe, D.B. (2017). Direct determination of diploid genome sequences. Genome Res 27(5), 757–767. doi: 10.1101/gr.214874.116.

Yamamoto, T. (1969). “Sex Differentiation,” in Fish Physiology, eds. W.S. Hoar & D.J. Randall. Academic Press), 117–175.

Yano, A., Guyomard, R., Nicol, B., Jouanno, E., Quillet, E., Klopp, C., et al. (2012). An immune-related gene evolved into the master sex-determining gene in rainbow trout, Oncorhynchus mykiss. Curr Biol 22(15), 1423–1428. doi: 10.1016/j.cub.2012.05.045.

Yasuda, S., Liu, C.C., Takahashi, S., Suiko, M., Chen, L., Snow, R., et al. (2005). Identification of a novel estrogen-sulfating cytosolic SULT from zebrafish: molecular cloning, expression, characterization, and ontogeny study. Biochem Biophys Res Commun 330(1), 219–225. doi: 10.1016/j.bbrc.2005.02.152.

Yazawa, R., Takeuchi, Y., Machida, Y., Amezawa, K., Kabeya, N., Tani, R., et al. (2019). Production of triploid eastern little tuna, Euthynnus affinis (Cantor, 1849). Aquaculture Research 50(5), 1422–1430. doi: 10.1111/are.14017.

