## Supplementary data for "Prediction of the sex-determination gene in tunas (*Thunnus* fishes)"

**Supplementary Fig. 1.** Comparison between the pseudo-haploid scaffolds obtained from Supernova2 and scaffolds M44/M175. The upper panel indicates the alignment between the Supernova2 scaffolds (scaffold IDs: 91818, 90643, 93129, and 275) and M44. The lower panel indicates the alignment between the Supernova2 scaffolds (scaffold IDs: 52814, 81188, 87065, 86196, 90712, 81419, 81140, 88535, 91688, and 82315) and M175. In the panels, SLR_M44_ and SLR_M175_ are highlighted in gray.

**Supplementary Fig. 2.** Mapped read depths around SLRs in 31 resequenced PBT samples. For each of the samples, SLR is highlighted in gray.

**Supplementary Fig. 3.** Genome-wide association with the sex of PBT using male masked reference sequences. Manhattan plot (left) and QQ plot (right) are shown. In the Manhattan plots, genome-wide significance cutoffs based on Bonferroni correction are indicate by a red line.

**Supplementary Fig. 4.** Comparison between male and female scaffolds encoding *sult1st6a*. The position of *sult1st6a* in scaffold M30 is indicated by an arrow.

**Supplementary Fig. 5.** Phylogenetic tree of *sult1st6* in fish species. For each of the nodes, the bootstrap probability is shown when it is >95%. The genes used in the main text (Fig. 9) are indicated with black circles. In each node, the Ensembl ID (e.g., ENSDARP00000023637 for zebrafish) denotes the protein sequence from which the CDS sequence is obtained.

**Supplementary Table 1. Protein-coding genes predicted in scaffold M175**

| No. | Start | End | Strand | Match in Ensembl | Identity (%) | Description |
| --- | --- | --- | --- | --- | --- | --- |
| g1 | 9390 | 9818 | + | ENSTRUP00000067783 | 41.3 | novel gene |
| g2 | 16389 | 17603 | - | ENSTRUP00000068256 | 63.7 | novel gene |
| g3 | 32614 | 64628 | + | ENSORLP00000007539 | 83.3 | estrogen sulfotransferase (*sult1st6*) |
| g4 | 76852 | 77963 | + | ENSTRUP00000072273 | 69.9 | novel gene |
| g5 | 82490 | 82919 | + | ENSTRUP00000067783 | 37.8 | novel gene |
| g6 | 118954 | 119841 | + | ENSORLP00000040498 | 83.8 | novel gene |
| g7 | 168127 | 168456 | + | ENSORLP00000044417 | 63.3 | novel gene |
| g8 | 176645 | 176974 | + | ENSORLP00000044417 | 63.3 | novel gene |
| g9 | 186243 | 186572 | + | ENSORLP00000044417 | 63.3 | novel gene |
| g10 | 195859 | 196188 | + | ENSORLP00000044417 | 62.4 | novel gene |
| g11 | 200940 | 201269 | + | ENSORLP00000044417 | 63.3 | novel gene |
| g12 | 207325 | 207654 | + | ENSORLP00000044417 | 65.1 | novel gene |

**Supplementary Fig. 1**

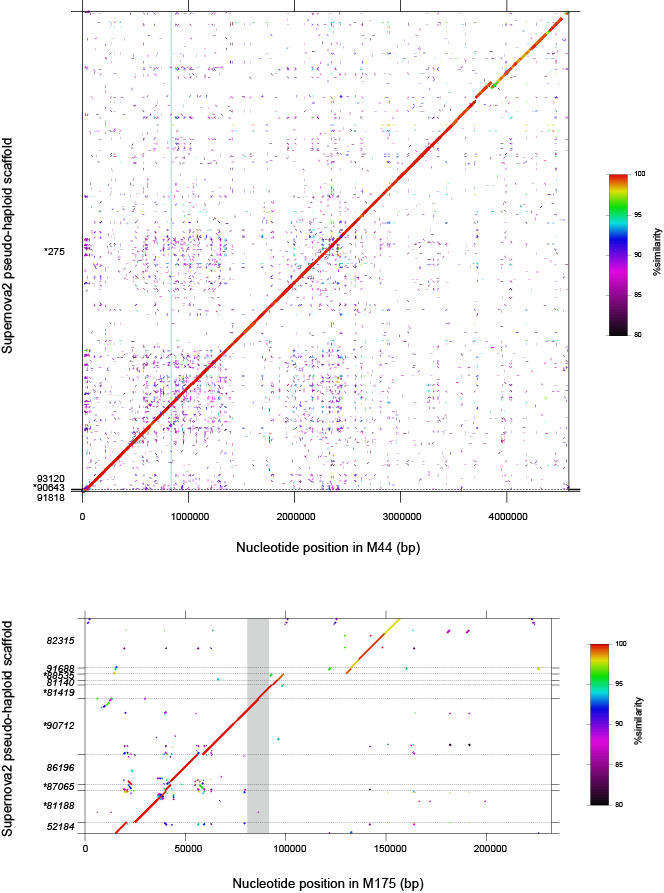

**Supplementary Fig. 2**

**Supplementary Fig. 3**

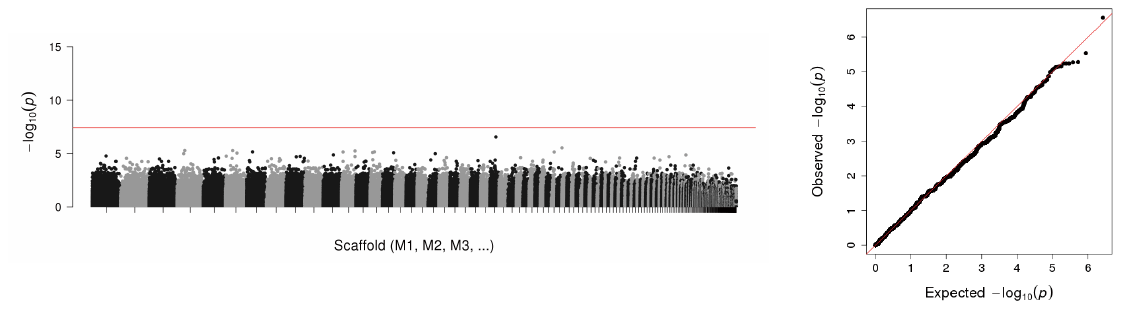

**Supplementary Fig. 4**

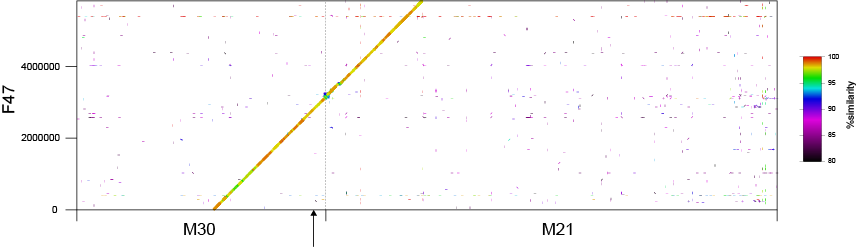

**Supplementary Fig. 5**

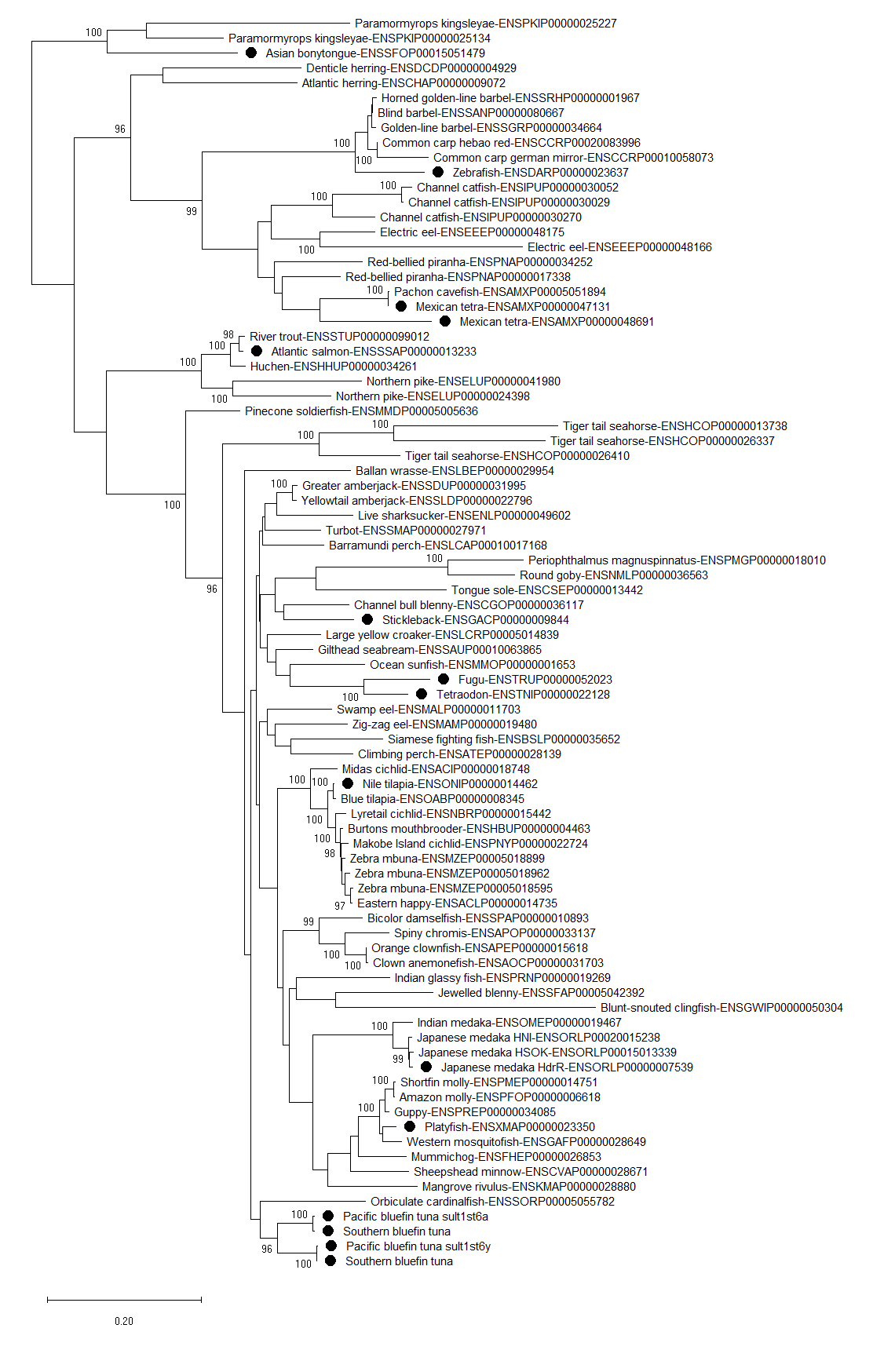
